# The challenge of structural heterogeneity in the native mass spectrometry studies of the SARS-CoV-2 spike protein interactions with its host cell-surface receptor

**DOI:** 10.1101/2021.06.20.449191

**Authors:** Yang Yang, Daniil G. Ivanov, Igor A. Kaltashov

**Affiliations:** Chemistry Department, University of Massachusetts-Amherst, Amherst, MA

## Abstract

Native mass spectrometry (MS) enjoyed tremendous success in the past two decades in a wide range of studies aiming at understanding the molecular mechanisms of physiological processes underlying a variety of pathologies and accelerating the drug discovery process. However, the success record of native MS has been surprisingly modest with respect to the most recent challenge facing the biomedical community – the novel coronavirus infection (COVID-19). The major reason for the paucity of successful studies that use native MS to target various aspects of SARS-CoV-2 interaction with its host is the extreme degree of structural heterogeneity of the viral protein playing a key role in the host cell invasion. Indeed, the SARS-CoV-2 spike protein (S-protein) is extensively glycosylated, presenting a formidable challenge for native mass spectrometry (MS) as a means of characterizing its interactions with both the host cell-surface receptor ACE2 and the drug candidates capable of disrupting this interaction. In this work we evaluate the utility of native MS complemented with the experimental methods using gas-phase chemistry (limited charge reduction) to obtain meaningful information on the association of the S1 domain of the S-protein with the ACE2 ectodomain, and the influence of a small synthetic heparinoid on this interaction. Native MS reveals the presence of several different S1 oligomers in solution and allows the stoichiometry of the most prominent S1/ACE2 complexes to be determined. This enables meaningful interpretation of the changes in native MS that are observed upon addition of a small synthetic heparinoid (the pentasaccharide fondaparinux) to the S1/ACE2 solution, confirming that the small polyanion destabilizes the protein/receptor binding.

## Introduction

The sudden emergence of the novel coronavirus infection^1^ (COVID-19) and it subsequent spread throughout the world has resulted in a truly unprecedented pandemic, which has already resulted in over 171 million confirmed cases and over 3.5 million deaths. Although the accelerated development and roll-out of several highly effective vaccines^2-6^ is likely to change the dynamic of this pandemic in the coming months, it currently shows no signs of abating world-wide, while the continuous and rapid evolution of the virus^7^ raises the specter of the possibility of the antibody immunity escape. The key element of the novel coronavirus SARS-CoV-2 allowing it to infect the host cells with high efficiency is the spike protein (S-protein). It enables the viral particle docking to the host cell via binding to its cell-surface receptor, a membrane-anchored angiotensin converting enzyme 2, ACE2^8^ (heparan sulfate, the part of the cell-surface proteoglycans, may assist in localizing SARS-CoV-2 near the surface of the epithelial cells, as is the case with a variety of other RNA viruses,^9-13^ but these interactions do not appear to play a significant role in SARS-CoV-2 cell adhesion^14^). Docking of the viral particle to the cell surface places the S-protein within the reach of the cell-surface proteases, several of which (*e*.*g*., furin and TMPRSS2) recognize the disordered segment connecting the S1 and S2 domains. Proteolytic processing of this segment (the so-called furin cleavage site) makes the fusion peptide available for anchoring into the cell membrane, followed by fusion of the virus with the cell^15^ (a mechanism common to all coronaviruses^16^). Inhibition of either of these processes should arrest the early stages of the cell infection cycle, placing the SARS-CoV-2 S-protein on the list of high-value therapeutic targets. This has fueled both extensive efforts to design novel drug candidates targeting the S-protein, and the relentless search for existing medicines capable of disrupting the S-protein interactions with its host cell-surface receptors. In the past, similar efforts were frequently aided by native mass spectrometry (native MS), a technique capable of detecting non-covalent complexes, such as multi-protein assemblies, as well as protein/small-molecule ligand associations under the physiologically relevant conditions.^17^ However, to date there had been very few reports in the literature describing successful application of native MS as a means of monitoring drug candidates’ interactions with the SARS-CoV-2 related therapeutic targets,^18,19^ despite the fact that multiple constructs representing various segments of the S-protein (including the entire ectodomain) are now commercially available from a range of sources.

The surprisingly modest success record of native MS when it comes to characterization of the interactions of the SARS-CoV-2 S-protein with its physiological targets and potential therapeutic agents is largely related to the extreme structural heterogeneity of this protein. Indeed, the polypeptide chain contains nearly two dozen N-glycosylation sites;^20-22^ and while the initial reports identified relatively few O-glycosylation sites,^22^ the latter have also been shown to be numerous in recent literature.^23^ The large number of the glycosylation sites and the microheterogeneity at each site give rise to a broad distribution of the total carbohydrate mass, which in the case of the trimeric S-protein was estimated to span a 100-200 kDa range.^24^ Such an enormous degree of structural heterogeneity is a serious challenge vis-à-vis extracting meaningful information on protein interactions from the native MS data. A strategy that is sometimes employed under similar circumstances aims at reducing the extent of glycosylation using either genetic tools (to eliminate at least some glycosylation sites from the polypeptide sequence^25^) or enzyme catalysis (to remove the N-glycans using enzymes from the PNGase family^26^). Unfortunately, the multiple glycans decorating the surface of the SARS-CoV-2 S-protein play an important role in both the viral entry into the host cell^27^ and the immune response evasion.^28^ Therefore, elimination of even a fraction of glycans (*e*.*g*., via de-*N*-glycosylation) is likely to introduce significant alterations in the S-protein interactions with its physiological partners, making the results of native MS measurements highly suspect.

A similar problem has been encountered in our recent studies of the receptor-binding domain (RBD) of the S-protein and its association with ACE2.^18^ Even though this 30 kDa segment of the S-protein contains only three N-glycosylation sites in addition to a few O-glycosylation sites, the extent of structural heterogeneity exhibited by the glycan chains is such that that the ionic signal in the ESI mass spectra of the S-protein RBD is presented by broad, poorly resolved distributions showing significant overlaps for different charge states. An attempt to mitigate this problem by switching to the glycan-free form of RBD (expressed in *E. coli*) failed due to the extreme instability of this form of the protein produced by prokaryotic systems. Eventually, meaningful information on RBD and its complexes with the cell-surface receptor was generated by supplementing the native MS measurements with the limited charge reduction.^18^ The latter technique isolates an ionic population within a narrow *m/z* window form convoluted ionic signal distributions in a mass spectrum, and subjects the isolated ions to interactions with either electrons or anions using the same set-up that is commonly utilized for wither electron capture- or electron transfer dissociation in commercial instruments.^29^ The resulting reduction of the number of positive charges carried by the polycationic species gives rise to a well-resolved charge ladder, from which both the charge (*z*) and the mass (*m*) of the precursor ions can be readily determined.^29^ While this approach was successful in providing interpretable data on the S-protein RBD/ACE2 interaction and the influence of short heparin oligomers on this complex stability, it is not clear whether larger segments of the S-protein and their complexes with the receptor will remain amenable to the native MS complemented by the limited charge reduction. In this work we consider the so-called S1 domain of the SARS-CoV-2 S-protein, a 100 kDa construct incorporating up to 37 possible glycosylation sites^23^ and its complexes with the receptor (human ACE2). In addition to the larger size and the significantly higher extent of glycosylation (compared to RBD), S1 also has a limited propensity to oligomerize (producing dimers and trimers, as well as high-molecular weight oligomers in solution). Nevertheless, a combination of the limited charge reduction and the data fitting procedures allows the S1/ACE2 complexes to de detected, and the influence of a short synthetic heparinoid (pentasaccharide fondaparinux) on the stability of this complex to be evaluated.

## Experimental

### Materials

The recombinant forms of human ACE2 ectodomain (residues 1-740) and the S1 segment of the SARS-CoV-2 S-protein (residues Val16-Arg685 with a C-terminal His-tag) expressed in a baculovirus system were purchased from Sino Biological (Wayne, PA). All proteins were extensively buffer exchanged in 150 mM NH_4_CH_3_CO_2_ prior to MS analyses (ultrafiltration at 3000 rpm with a 10 kDa MW cut-off filters for 40 min repeated 3 times). Fondaparinux was purchased from Sigma-Aldrich (St. Louis, MO). All solvents and buffers were of analytical grade or higher.

### Methods

Native MS measurements were carried out using a Synapt G2-Si (Waters, Milford, MA) hybrid quadrupole/time-of-flight mass spectrometer equipped with a nanospray ion source. The following ion source parameters were used to ensure stability of the non-covalent complexes in the gas phase: capillary voltage, 1.5 kV; sampling cone voltage, 80 V; source offset, 80 V; trap CE, 4 V; trap DC bias, 3 V; and transfer CE, 0 V. Ion selection for limited charge reduction was achieved by setting the appropriate quadrupole selection parameters (LM resolution set at 4.5). Limited charge reduction was initiated by allowing the m/z-selected multiply charged ions to interact with 1,3-dicyanobenzene anions for 0.6 msec after setting the trap wave height as 0.3 V and optimizing the discharge current. All MS measurements were repeated at least twice to ensure consistency of the results. Fitting of the experimental data was performed using Anaconda distribution of Python 3. The methods implemented in SciPy library from the distribution were used to perform fitting of normal distributions to experimental data by Levenberg-Marquardt algorithm, as well as to predict the intensities of charge states that lie under the baseline assuming a normal distribution of charges between different charge states in native spray.

## Results and Discussion

The ESI mass spectrum of the S1 domain of the SARS-CoV-2 S-protein acquired under near-native conditions in solution displays a convoluted, nearly-continuum distribution of the ionic signal (**Figure 1**). Despite the highly convoluted appearance, several features appear to be discernable in the spectrum, such as the ionic signal confined to the m/z ranges 5,000-6,500; 7,000-8,000; 8,000-11,000 and 13,500-20,000. The lowest-m/z spectral feature appears to contain partially resolved charge states, while the rest display a continuum signal. Performing limited charge reduction measurements on the lowest-m/z spectral feature (the blue trace in **Figure 1**) confirms that the poorly resolved peaks within this ionic signal cluster correspond to individual charge states of a macromolecular species with an average mass of 92.8±3.4 kDa. The mass of the recombinant S1 construct used in this work is 76.7 kDa excluding all glycans. In order to evaluate the total glycan mass, we used the masses of the most abundant glycoforms within the S1 domain, as reported by Shajahan et al.,^22^ and calculated a net mass of glycans as 14.1 kDa, leading to identification of the lowest-m/z feature as the S1 monomer. Processing of the second cluster of the ionic signal (m/z 7,000-8,000) using limited charge reduction of the ionic population at the apex of the signal distribution also gave rise to a well-resolved charge ladder (the purple trace in **Figure 1**). Based on this charge ladder, the mass of the most abundant species giving rise to the ionic signal within m/z region 7,000-8,000 was calculated as 184.8±4.8 kDa, allowing it to be identified as S1 dimer.

**Figure 1.**
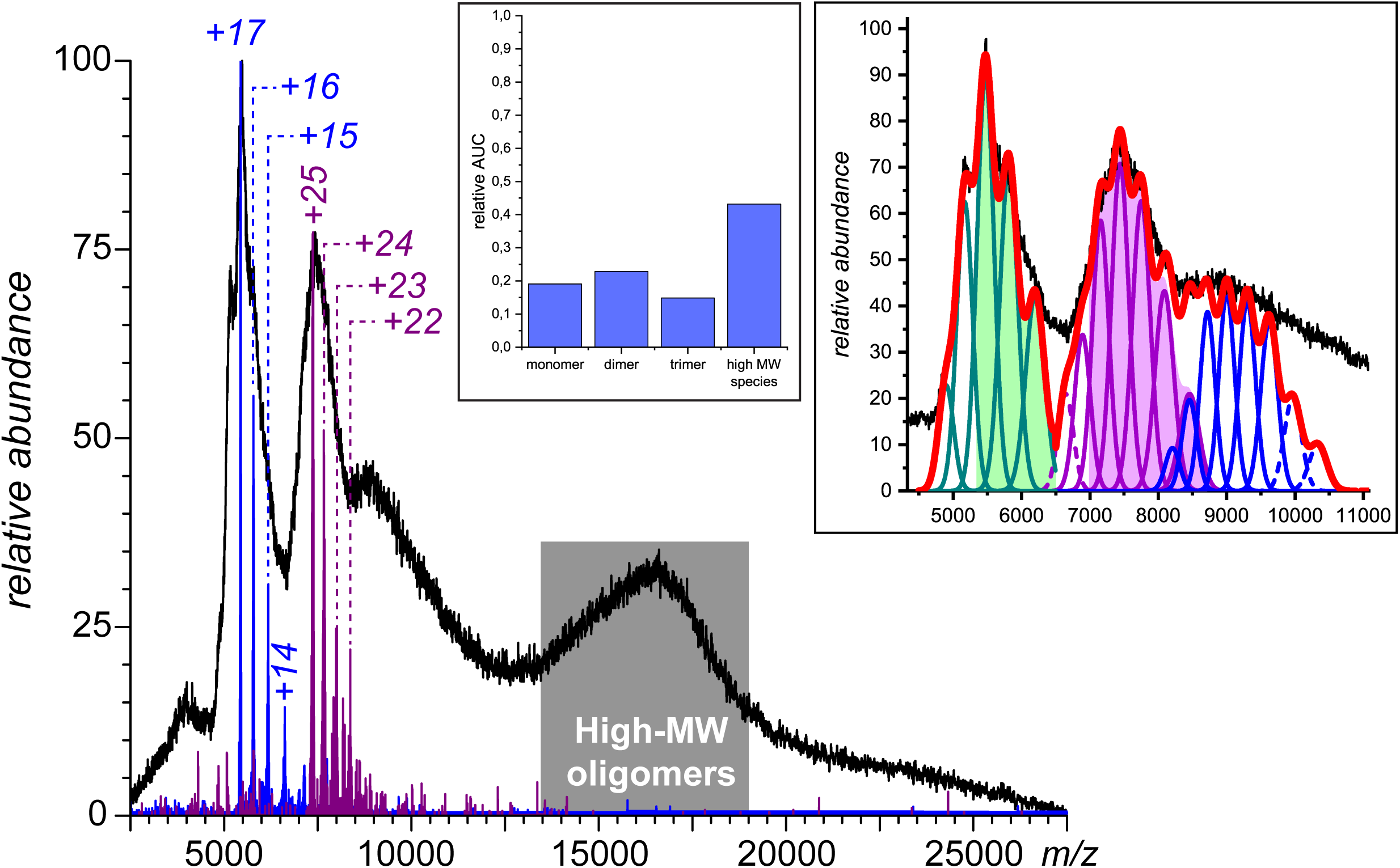
An ESI mass spectrum of S1 (0.28 mg/ml in 150 mM aqueous ammonium acetate) shown as a black trace, and mass spectra obtained by limited charge reduction of ionic populations representing the highest-intensity signal in two most abundant spectral features (blue and purple). The two charge ladders generated by the limited charge reduction allow the two spectral features to be identified as ions corresponding to S1 monomer (measured mass for the most abundant ionic species 92.8 kDa), S2 dimer (measured mass for the most abundant ionic species 184.8 kDa). The inset shows abundance distribution of various oligomeric states of S1 calculated on the basis of their molar fractions.

Identification of the two main spectral features in the mass spectrum shown in **Figure 1** as the monomeric and dimeric forms of S1 allowed us to test another approach to interpretation of unresolved MS data, which is based on fitting the experimental data with the calculated ionic profiles. Specifically, the charge state assignments provided by the limited charge reduction work on the monomeric S1 signal were used to generate the best fit for the segment of the mass spectrum confined to the m/z range 5,000-6,500 (**Figure 1a**). The mass profile of the S1 monomer yielding the best fit with the experimental data (the least deviation in Euclidean distance between experimental profile and the approximated one was used as quality criteria in all approximations) was then used to calculate the ionic signal profile for the S2 dimer in the m/z region 7,000-8,000 relying on the charge assignments generated by the limited charge reduction work. While the mass distribution of the S1 dimers (produced by convolution of the mass profile for the S1 monomers) and the charge states were fixed, the relative abundance of the ionic signal at each charge state was allowed to vary to obtain the best fit with the experimental data. The resulting profile is presented as an inset in the **Figure 1**. The extension of this procedure to the third abundant spectral feature (m/z range 8,000-11,000) and assuming that it represents the trimeric state of S1 allows a reasonable fit with the experimental data to be obtained using charge states +27 through +34. Although the latter relies exclusively on the fitting procedure (no limited charge reduction data were available for this cluster of ionic signal due to the high ionic m/z values and the low signal abundance), indirect verification can be carried out by comparing the extent of multiple charging of all three states of S1 in **Figure 1** and relating these numbers to the known empirical relationship between the average ionic charge (Z_av._) and the solvent-exposed surface area of the corresponding species in solution (S):^30,31^

**Figure 1a.** An approximation of ESI spectra by combination of convolved normal distributions of pure S1 monomer compounds with different charge states. Green, pink and blue area refer to approximated S1 monomer, dimer and trimer signals. The solid lines green, pink and blue lines refer to those charge state signals whose intensity was approximated by fitting of experimental intensity. The dashed lines refer to those charge state signal, which lie under the baseline signal, and so, were approximated by fitting to normal distribution defined from more intense approximated charge states according to [30]. The red line shows the signal approximation.

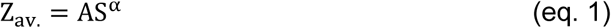

where A is a constant and the empirical value of α is 0.69±0.02. Although the surface area values are usually calculated based on the protein crystal structure, using the published crystal structures of the SARS-CoV-2 S-protein to calculate the surface areas of S1 monomers, dimers and trimers by simply truncating the protein sequence and removing irrelevant segments is likely to generate misleading results for two reasons. First, it is not clear whether the monomeric and/or dimeric S1 would maintain the same conformation as the trimeric form of this protein domain. Second, the intrinsic flexibility of the glycan chains makes them mostly invisible in crystal structures of glycoproteins; as a result, their contributions to the overall surface area of S1 (which is extensively glycosylated) will not be accounted for. Therefore, instead of using the available crystal structures of the S-protein, we obtained rough estimates of expected ratios of the average charges for the three different oligomeric states of S1 (monomer, dimer and trimer) using a quasi-spherical approximation. Each protein is assumed to have a spherical shape and density that is uniform across all three forms of S1, i.e. the mass of each species has a straightforward relationship with its dimensions in solution:

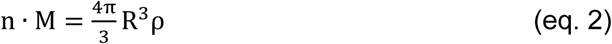

where n is the number of monomeric units in the assembly, M is the monomer mass, R is the radius of the protein particle in spherical approximation, and ρ is its density. In this approximation the surface area of the protein particle can be expressed as a function of its mass simply as

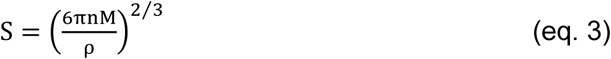

and the expected average number of charges accumulated by the corresponding ESI-generated ions would be

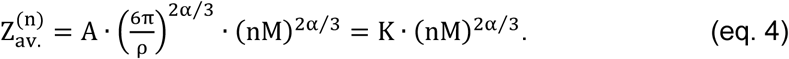

According to this expression, the ratio of the average charge states of ions representing dimeric (n = 2) and monomeric (n = 1) forms of S1 should be 1.38, a value that is remarkably close to the ratio of the most abundant charge states for these two groups of ions in the mass spectrum shown in **Figure 1** (25/17 = 1.47). Likewise, the ratio of the average charge states for trimeric and dimeric forms of S1 estimated using (eq. 4) is 1.21, which is in excellent agreement with the ratio of the charge states obtained using charge states of the most abundant ions in each cluster (31/25 = 1.24).

A unique feature of the mass spectrum presented in **Figure 1** is the ionic signal dominating the high m/z region of the spectrum. Prior to assigning this feature at m/z > 13,500 as high-molecular weight solution-borne aggregates of S1, one alternative explanation must be considered, namely the gas-phase dissociation of S1 trimers (and, possibly, dimers) proceeding via the asymmetric charge partitioning.^32-34^ The latter phenomenon results in ejection of a high-charge density polypeptide chain from a protein complex in the gas phase, leaving behind the remaining part of the complex carrying a disproportionately low number of charges (which appear in the mass spectra at high m/z values). The resulting spectral features can lead to misinterpretation of the MS data; therefore, the possibility of the occurrence of the asymmetric charge partitioning must be considered when the ionic signal in mass spectra appears highly convoluted and populates the high m/z region.^35^ For example, one might argue that the high-m/z spectral feature in **Figure 1** presents the low-charge density fragments generated by dissociation of the trimeric S1 species. However, a putative asymmetric charge partitioning affecting the (S1)_3_^31+^ species and leading to ejection of (S1)^17+^ chains would give rise to (S1)_2_^14+^ ions, which would appear at m/z 13,000 (a few thousand m/z units away from the apex of the experimentally observed ionic distribution). Likewise, a putative dimer-to-monomer dissociation is expected to generate a low-charge density fragment at m/z 11,500, which is even further away from the apex of the experimentally observed ionic signal in the high-m/z region of the mass spectrum (m/z 16,500). Therefore, it appears unlikely that the spectral feature dominating the high m/z region of the mass spectrum shown in **Figure 1** represents products of the gas-phase dissociation, and protein aggregation in solution is the only process that can be invoked as a means of explaining the origins of this ionic signal. The average size of these high-MW oligomers of S1 can be estimated using an approach similar to that described earlier in this paragraph (see eqs. 1-4). Indeed, expression (eq. 4) can be re-written as

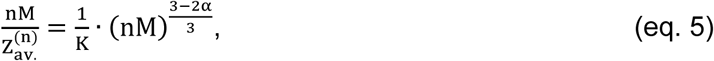

and equating the (nM)/Z_av._ ratio to an m/z value of the corresponding oligomer, one can present this expression in the following simplified form:

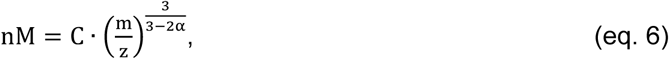

where C is a constant. This allows the number of monomeric units of S1 incorporated in the high-MW oligomers giving rise to ions at m/z 16,500 (the apex of the high m/z spectral feature) to be estimated as n ≈ 9 (using the m/z value of 7,400 for the dimeric S1). While the relative abundance of these aggregates appears to be lowest among all oligomeric states of S1 giving rise to the four distinct spectral features visible in **Figure 1**, one needs to remember that MS signal reflects molar concentrations of analytes, rather than their weight concentrations.^36^ In other words, in the absence of any biases due to ionization and ion transmission differences, a single S1 monomer would give rise to an ionic signal of the same intensity as a high-MW aggregate (e.g., S1_9_), even though conventional detection by UV would indicate a 1:9 signal ratio. Converting the MS signal abundance ratio to fractional mass concentrations presents a more realistic picture of the protein material distribution among all detected oligomeric states, with the high-MW aggregate being the most abundant species and accounting for a nearly 43% weight fraction of the total protein.

The ectodomain of ACE2, the binding target of the SARS-CoV-2 on the host cell surface, exists as a non-covalent dimer in solution,^18^ and adding this protein to the S1 solution results in a dramatic change of the mass spectra (**Figure 2**). The ACE2/S1 mass spectra exhibit a strong dependence on the stoichiometric ratio of the two proteins in solution. For example, molar excess of ACE2 over S1 (corresponding to the *ca*. 3.75:3 dimeric ACE2:monomeric S1 ratio) gives rise to a distinct cluster of peaks in the mass spectrum in the *m/z* region 8,000-9,000 with discernable charge states +32 through +34, and a mass consistent with a single S1 monomeric unit bound to an ACE2 receptor. An increase of the S1 content gives rise to another distinct spectral feature (confined to the *m/z* range 9,000-10,500), which becomes a dominant feature at the ACE2/S1 ratio of 1:3 (see the purple trace in **Figure 2**). The strong dependence of the relative abundance of these ions on the ACE2/S1 ratio suggests that they represent an ACE2/S1 complex with two monomeric S1 subunits associated with a single ACE2 dimer. Indeed, using the spectral convolution/data fitting procedure similar to that applied earlier to model the appearance of S1 dimers and trimers (*vide supra*), we were able to assign this spectral feature as a cluster of ions with an average mass of 357 kDa (ACE2 dimer and two monomeric S1 units) with charge states ranging from +35 to +47.

**Figure 2.**
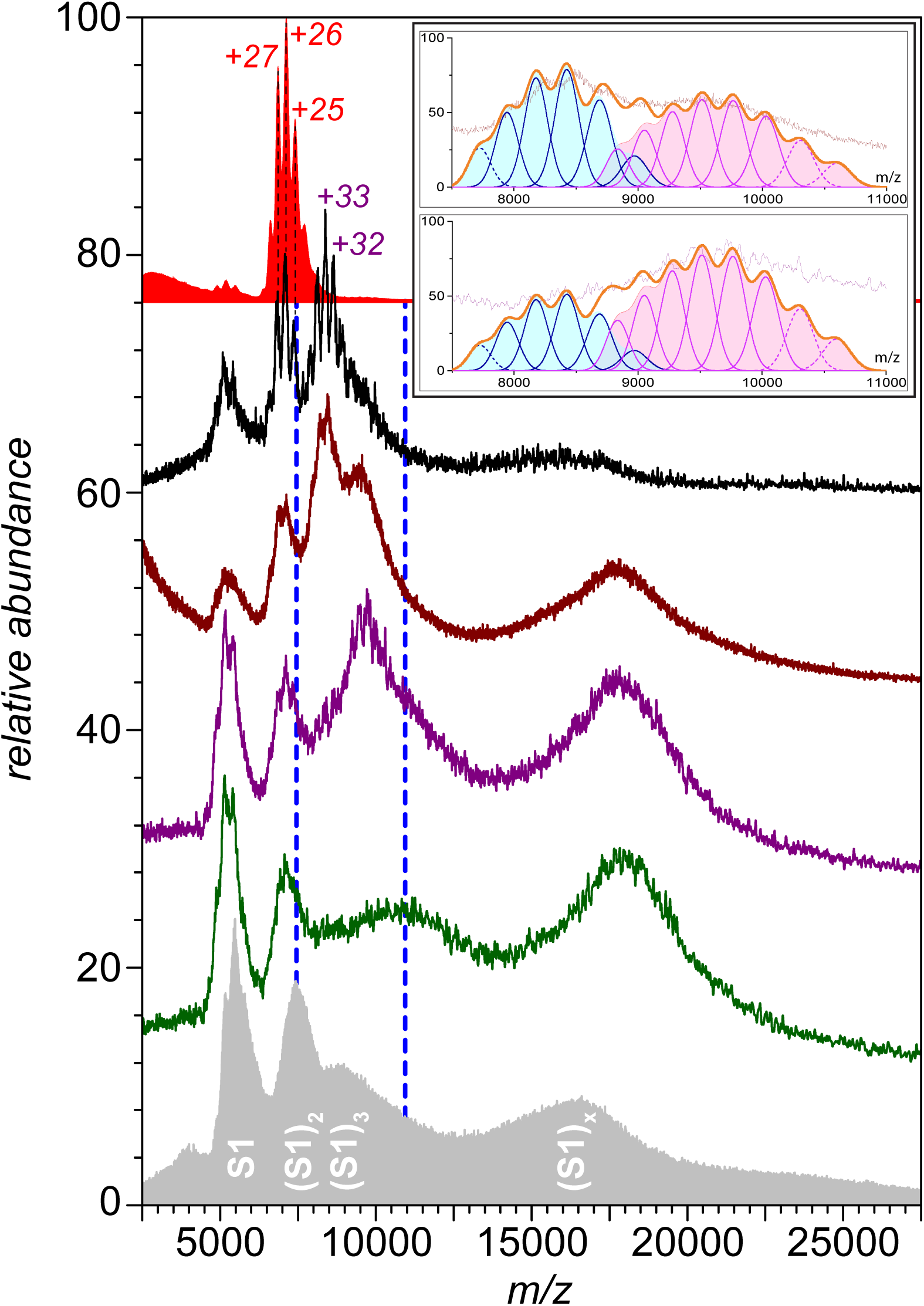
An ESI mass spectra of ACE2 (top), ACE2/S1 mixtures at 3.75:3, 2.5:3, 1:3 and 0.5:3 (traces 2-5 from the top, respectively), and a reference spectrum of S1 (bottom). The inset shows the approximation (that was done as in Figure 1a) of charge states of [S1+ACE2] (pink) and [2 S1+ACE2] (blue) compounds for 2.5:3, 1:3 mixtures.

Two other spectral features that gain prominence as the molar ratio of S1 continues to increase are the signals representing the free (unbound) S1 monomers and dimers (*m/z* 4,500-8,000) and the high-molecular weight species (*m/z* above 15,000). Importantly, the latter is shifted upwards on the *m/z* scale with respect to the high-MW species observed in the mass spectrum of S1 alone (**Figure 1**), suggesting that the high-MW S1 aggregates (previously identified as nonamers, *vide supra*) can associate with ACE2. The mass increase associated with this *m/z* shift of the ionic signal (from *ca*. 16,500 to *ca*. 18,100) can be estimated using the quasi-spherical approximation presented above and (eq. 6), providing a *ca*. 160 kDa increment, a value that is within 15% of the average mass of an ACE2 dimer. Remarkably, this increment appears to be nearly constant across a wide range of ACE2/S1 ratios (see **Figure 2**), in contrast to the evolution of the ionic signal in the 11,000-13,000 *m/z* region. The monotonic increase of the latter upon the increase of the S1/ACE2 ration likely indicates the appearance of ionic species representing a complex, in which one of the monomeric subunits a single ACE2 dimer is associated with S1 monomer, and the other is bound to the (S1)_2_ dimer. A more definitive assignment of the stoichiometries of various S1/ACE2 complexes observed in ESI mass spectra shown in **Figure 2** would require the use of limited charge reduction of ions in the gas phase preceded by their supercharging in solution to shift the ionic signal to m/z region where the precursor mass selection can be made prior to initiating the charge-exchange reactions in the gas phase.^37^

While the analysis of the S1/ACE2 binding presented in the preceding paragraph indicates that the majority of the spectral features can be assigned, it is not clear whether the level of resolution attained in these measurements would be sufficient for evaluation of the influence of various therapeutic agents on the stability of the S1/ACE2 association. Previously we demonstrated that even relatively short heparin oligomers can disrupt the interaction between the S-protein RBD and ACE2.^18^ Specifically, a short (pentasaccharide) synthetic heparin analog fondaparinux was shown to be able to associate with RBD and induce conformational changes that allosterically disrupted the RBD/ACE2 binding.^18^ Unfortunately, the fondaparinux mass is too small compared to the mass variance of S1 (1728 Da vs. 3.4 kDa), making it nearly impossible to detect association of a single fondaparinux molecule with S1. Nevertheless, the effect of this short heparinoid on the S1/ACE2 interaction can be readily assessed using native MS. Indeed, addition of fondaparinux to the S1/ACE2 solution (to a final concentration of 1.16 μM, which is a ca. 1.89 molar excess over S1) results in a dramatic change of the mass spectrum of the 3:2.5 S1/ACE2 mixture (**Figure 3**). Specifically, the ionic signal corresponding to the free ACE2 dimers in solution becomes the dominant feature of the mass spectrum, while the relative abundance of ions representing the ACE2·(S1)_2_ complexes suffers a dramatic decrease. In fact, the latter is no longer a distinct, second-most-abundant spectral feature, but rather has an appearance of a high-*m/z* shoulder of the spectral feature corresponding to the ACE2·S1 complexes (which is also greatly reduced compared to the ACE2 ionic signal in the presence of fondaparinux).

**Figure 3.**
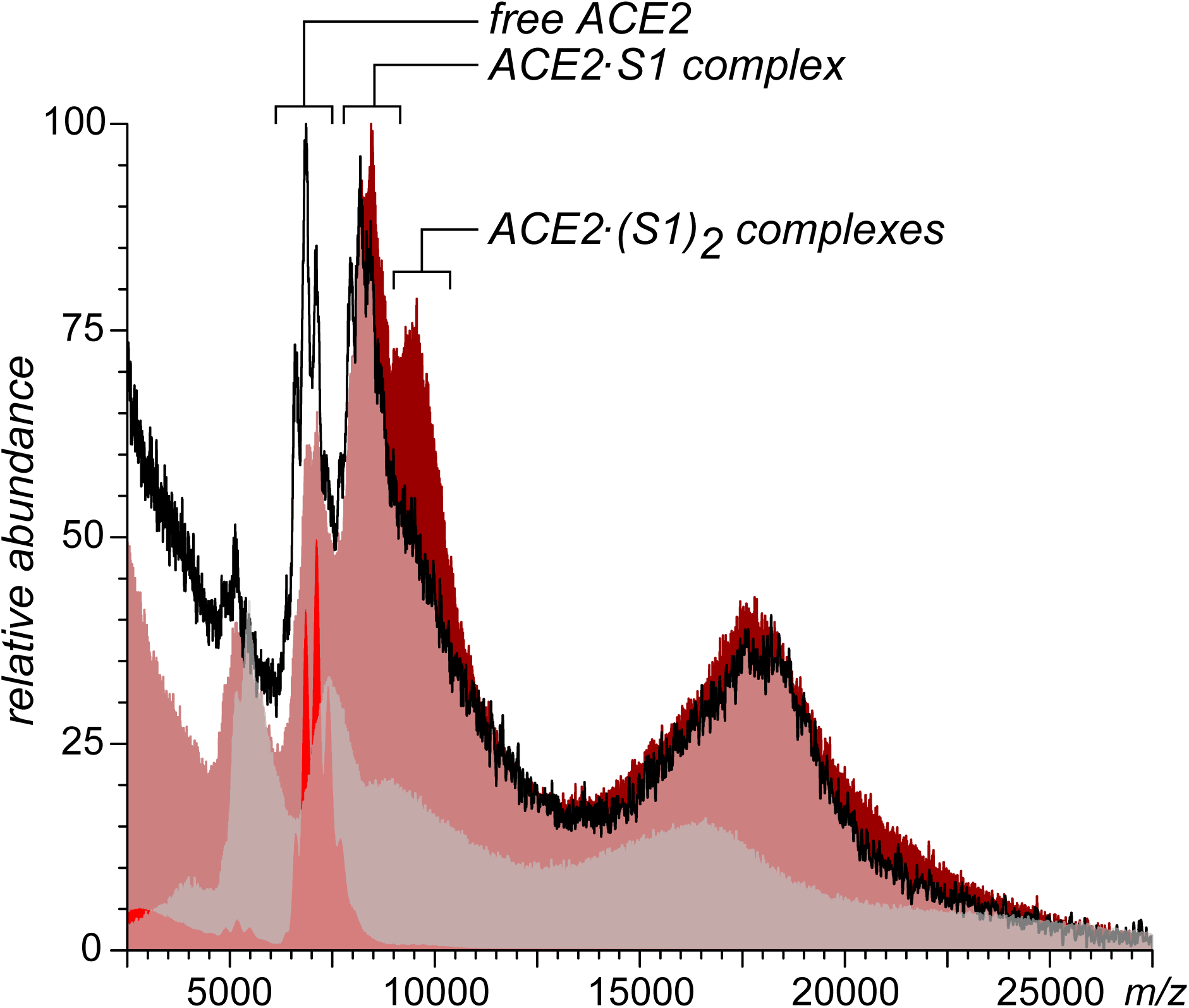
ESI mass spectra of the ACE2/S1 mixture (2.5:3) before and after addition of 1.16 μM fondaparinux shown as the brown and black traces, respectively. The reference mass spectra of S1 and ACE2 are shown in gray and red (not to scale).

Careful examination of the high-*m/z* region of the mass spectrum of the ACE2/S1 mixture acquired in the presence of fondaparinux (**Figure 3**) reveals that the spectral feature previously assigned as a complex formed by association of a single ACE2 dimer with a high-MW S1 aggregate does not appear to be affected by the small heparinoid. Indeed, the position of the apex of this spectral feature appears to be nearly identical to that observed in the mass spectrum of the ACE2/S1 mixture in the absence of fondaparinux, and exceeding the apex of the ionic signal representing the ACE2-free S1 aggregates (as seen in the mass spectrum of S1) by over 1,500 *m/z* units. This apparent (and certainly surprising) insensitivity of the high-MW S1 aggregates to fondaparinux vis-à-vis their ability to associate with ACE2 in solution likely points out to the fact that the disruption of the RBD/ACE2 association is allosteric in nature and is triggered by the heparinoid binding to the positive patch on the RBD surface that is distal to the receptor binding site, as suggested by the previous molecular modeling studies.^18^ The access to this distal site is likely to be obstructed in the high-MW S1 aggregates, preventing fondaparinux binding and the ensuing conformational changes. It is also possible that the allosteric conformational changes that destabilize the ACE2 binding site on the RBD surface are at least partially inhibited within the high-MW S1 aggregates due to the tight packing of the monomeric S1 subunits.

Although the dramatic changes in the appearance of the mass spectra of the ACE2/S1 mixture in the presence of fondaparinux do not provide quantitative information on the receptor affinity changes of S1 in the presence of the small heparinoid, they are sufficient to make a definitive qualitative conclusion that the latter is an effective disruptor of the ACE2/S1 interaction. Furthermore, the apparent inability of fondaparinux to induce dissociation of ACE2 from the high-MW S1 aggregates provides additional support for the previously formulated hypothesis that heparinoids disrupt the S-protein binding to its cell-surface receptor ACE2 allosterically via binding to a site that is distal to the ACE2 interface. Therefore, despite the lack of resolution required for straightforward identification of all ions produced upon ionization of the S1 domain alone and its mixture with ACE2, native MS of these highly heterogeneous systems not only provides invaluable information on the oligomerization state of S1 and their binding to ACE2, but also highlights the ability of short heparin oligomers to disrupt most (but not all) such interactions. Therefore, native MS is already capable of providing important mechanistic insights into the interactions of large components of the SARS-CoV-2 S-protein with therapeutic agents, and their influence on the S-protein interaction with its physiological targets. Broader utilization of native MS to support drug repurposing efforts will further catalyze the progress towards developing a broad spectrum of viable therapeutic option for both prevention and treatment of COVID-19.

## Conclusions

Despite the herculean efforts of the entire biomedical community, and the significant resources devoted to the task of finding effective treatments for the novel coronavirus infection (SARS-CoV-2), the therapeutic options remain disappointingly limited. While the large-scale vaccination campaigns are expected to tame the COVID-19 pandemic in the coming months, the lack of international coordination/collaboration and the alarmingly rapid emergence of novel strains^38^ are likely to present a significant challenge to these efforts due to the high transmissibility of many novel variants and the reduced vaccine effectiveness against them.^39,40^ The vulnerability of the highly specific medicines, such as mAbs, to the SARS-CoV-2 mutations places a premium on the on-going efforts to develop broad-spectrum therapeutics (including both novel and repurposed drugs) that would remain effective not only against the ever increasing range of SARS-CoV-2 variants, but also the future threats posed by the possible introduction of other coronavidæ to the human population via either zoonotic or lab escape routes.^41^ Native MS has been used as an effective analytical tool in a wide range of drug discovery/development efforts, but it remains disappointingly underutilized in the studies aiming at developing novel therapeutics that target SARS-CoV-2. The large size and the extreme structural heterogeneity of the SARS-CoV-2 S-protein, the major determinant of its infectivity (and the prime target of both vaccine and drug development efforts), are the two major factors limiting the utility of native MS in this field. Our work with the S1 domain of the S-protein demonstrates that even if the desired level of resolution is unattainable, native MS provide valuable information on the interaction between this protein and its physiological receptor. Furthermore, native MS allows the influence of small-molecule drugs in such interactions to be evaluated, arguing for a broader utilization of this technique in the screening efforts. The on-going efforts to expand the capabilities of native MS in the realm of large and structurally heterogeneous targets (such as the development of the limited charge reduction technique,^29^ which already demonstrated its effectiveness in targeting highly heterogeneous systems ranging from PEGylated^42^ and heavily haptenated^43^ proteins to unfractionated heparin^44^) will undoubtedly expand both the range of molecular targets and provide further impetus to the search for effective and safe medicines against the novel coronavirus.

## Authors’ contributions

YY and IK designed the study and planned the experimental work; YY carried out the experimental work; DI designed the data fitting algorithm; YY and DI processed the experimental data; IK wrote the manuscript. All authors gave their approval to the submitted version of the manuscript.

## Acknowledgements

This work was supported by a grant R01 GM112666 from the National Institutes of Health. All MS measurements were carried out in the Mass Spectrometry Core Facility at UMass-Amherst.

